# A modular platform for scalable recombinant production of highly toxic bacterial proteins

**DOI:** 10.64898/2026.07.31.742022

**Authors:** Rina Fraenkel, Inbar Cahana, Tomer Sivan, Tal Fisher, Hen Nadav, Shani Bruchim, Noam Deouell, Shani Cheskis, Maor Shalom, Lionel Imbert, Asaf Levy, Netanel Tzarum

## Abstract

Bacterial protein toxins constitute a vast and largely untapped reservoir of antimicrobial activities with substantial therapeutic and biotechnological potential. However, their intrinsic toxicity frequently prevents stable recombinant expression in bacterial hosts, creating a major bottleneck for biochemical characterization, structural analysis, and development as antimicrobial agents. Here, we present a modular platform for the scalable recombinant production of highly toxic bacterial proteins based on transient intramolecular toxin neutralization. The strategy covalently links each toxin to its cognate immunity protein, promoting neutralization during biosynthesis while permitting recovery of the native toxin through site-specific proteolytic cleavage. Using this approach, we produced multiple previously intractable polymorphic toxin domains that could not be obtained using conventional inducible expression, toxin–immunity co-expression, or bacterial cell-free systems. We further streamlined the production workflow through intracellular protease-mediated cleavage, reducing the purification process from four steps to two and increasing protein recovery. To address cases in which native immunity proteins were insufficient, we incorporated computational protein design to engineer improved toxin-binding partners, enabling production of an additional toxin that remained refractory to the original platform. Purified toxins retained enzymatic activity following denaturation and refolding, confirming recovery of functional proteins and enabling identification of a previously uncharacterized nuclease activity. Together, these findings establish a scalable and adaptable microbial biotechnology platform for the production of intrinsically toxic proteins. The integration of transient intramolecular neutralization with computational engineering provides a route toward systematic production and characterization of toxic proteins for antimicrobial discovery, structural biology, protein engineering, and future biotechnological applications.

## Introduction

Antimicrobial-resistant (AMR) bacterial pathogens caused an estimated 4.95 million deaths worldwide in 2019, including 1.27 million deaths directly attributed to bacterial AMR (Antimicrobial Resistance, 2022). In parallel, invasive fungal diseases represent an additional and growing global health burden, causing approximately 6.5 million infections and 3.8 million deaths annually, with 2.5 million deaths directly attributable to fungal infections (Denning, 2024). The limited number of antifungal drug classes, together with the rapid emergence of drug-resistant fungal strains, further exacerbates this crisis. Without effective intervention, AMR-associated mortality is projected to reach 10 million deaths annually by 2050, prompting the World Health Organization (WHO) to designate AMR as one of the top global public health threats (WHO fungal priority pathogens list to guide research, development, and public health action, 2022).

The rapid global spread of multidrug-resistant bacterial and fungal pathogens underscores the urgent need for antimicrobial agents with novel mechanisms of action (MoAs) that are less likely to resistance. However, discovering and developing new antimicrobial compounds face major scientific, economic, and regulatory challenges, resulting in a diminishing clinical pipeline (Gigante, et al., 2024). Antifungal drug development is particularly challenging because fungal pathogens, as eukaryotes, are closely related to humans, limiting the availability of selective therapeutic targets and increasing the risk of host toxicity (Roemer and Krysan, 2014, Roy, et al., 2023, Li, et al., 2025, Souza, et al., 2025). These limitations highlight the importance of exploring alternative classes of antimicrobial agents with distinct biological activities and mechanisms.

Among the most promising alternatives to conventional antibiotics are naturally occurring antimicrobial proteins, a diverse class of molecules that mediate microbial competition across all domains of life. These proteins employ a wide range of mechanisms to eliminate competing microorganisms, including disruption of membranes, inhibition of cell-wall biosynthesis, interference with protein synthesis, and degradation of nucleic acids (Zhang and Gallo, 2016, Lei, et al., 2019, Mookherjee, et al., 2020). Their therapeutic potential is exemplified by the clinical success of peptide- and protein-based antimicrobials such as polymyxins, gramicidin, nisin, and echinocandins (Guiotto, et al., 2003, Zheng, et al., 2025). Among these molecules, antimicrobial small proteins, typically 50-150 amino acids in length, represent a particularly underexplored class. Many antimicrobial small proteins function as compact enzymatic toxins that disable essential cellular processes with remarkable potency. A prominent example is the bacterial polymorphic toxin system, which consists of modular proteins carrying highly diverse toxic C-terminal effector domains that target essential cellular processes, including nucleic acid metabolism, protein synthesis, and cell-wall biogenesis (Zhang, et al., 2012, Jamet and Nassif, 2015, Ruhe, et al., 2020). To prevent self-intoxication, toxin-producing bacteria co-express highly specific cognate immunity proteins that bind and neutralize their corresponding toxin domains. Their exceptional potency, mechanistic diversity, and modular organization make polymorphic toxins attractive candidates for next-generation antimicrobial therapeutics and valuable tools for discovering previously unknown antibacterial mechanisms.

Despite their considerable biological and therapeutic potential, recombinant production of enzymatic antimicrobial toxins remains a major unsolved challenge. Recombinant protein production is routinely performed in *Escherichia coli* (*E. coli*) because of its rapid growth, low cost, and scalability (Rosano and Ceccarelli, 2014). However, unlike most recombinant proteins, antimicrobial toxins directly attack highly conserved bacterial processes required for protein synthesis and cell viability (Baneyx, 1999). Consequently, even low levels of basal expression frequently impose strong selective pressure for plasmid loss or inactivating mutations while simultaneously impairing protein accumulation through growth inhibition, proteolytic degradation, aggregation, and inclusion-body formation (Sorensen and Mortensen, 2005, Rosano and Ceccarelli, 2014, Hagan, et al., 2023). Existing recombinant production strategies—including tightly regulated promoters, co-expression of immunity proteins, inclusion-body purification, and cell-free expression—are often protein-specific, labor-intensive, costly, or difficult to scale.As a result, recombinant production remains the principal bottleneck preventing systematic biochemical, structural, and mechanistic investigation of these proteins.

An additional essential step in developing antimicrobial toxins into effective therapeutics is understanding their MoAs. Mechanistic characterization elucidates the structural and functional determinants that govern potency, specificity, and stability, and enables rational engineering to improve efficacy and minimize the emergence of resistance (Raheem and Straus, 2019, Li, et al., 2022). Nevertheless, many toxin proteins remain mechanistically uncharacterized, representing a vast, largely untapped reservoir of novel antimicrobial activities. Systematic discovery of novel toxin proteins and their unique MoAs could uncover fundamentally new strategies to combat bacterial and fungal pathogens and overcome existing resistance mechanisms (Nachmias, et al., 2024). Realizing the therapeutic and biotechnological potential of these highly toxic proteins critically depends on scalable, tightly controlled bioproduction platforms that enable efficient recombinant expression and manufacturing.

Recent advances in genomics and high-throughput bioinformatics provide unprecedented opportunities to systematically identify previously unrecognized antimicrobial toxin proteins and investigate their biological activities. Leveraging these approaches, we developed a computational framework to identify novel antimicrobial toxin proteins and applied it to analyze more than 100,000 bacterial genomes (Nachmias, et al., 2024). This effort led to the discovery of nine previously uncharacterized toxin domains within polymorphic toxin systems (PT1-PT9), each representing a distinct protein family defined by sequence similarity (Zhang, et al., 2012, Ruhe, et al., 2020). For a subset of PTs, we identified their associated immunity genes (Polymorphic Immunity genes, PIMs) that protect the toxin-producing microorganism from self-toxicity (Aoki, et al., 2010). Functional validation confirmed strong antibacterial activity against E. coli and, for several PTs, demonstrated strong antifungal activity against multiple fungal pathogens, including top-priority human pathogens listed by the WHO.

The identification of these antimicrobial toxins created an opportunity to explore a new class of antimicrobial proteins. However, realizing their scientific and translational potential requires scalable production of purified, biologically active proteins. Because conventional recombinant expression repeatedly failed for these highly toxic proteins (Nachmias, et al., 2024), developing a robust, scalable, and broadly applicable production platform became the critical next step toward their biochemical characterization and future biotechnological exploitation.

Here, we address this challenge by establishing a modular production platform based on transient intramolecular neutralization of toxin activity during recombinant expression. By integrating toxin-immunity fusion proteins, streamlined purification, and computational protein engineering, our approach overcomes key bottlenecks in the recombinant production of highly toxic proteins. We demonstrate the platform using multiple polymorphic toxin effector domains that were previously refractory to recombinant expression and show that the recovered proteins retain their native enzymatic activity. Collectively, this work establishes a scalable microbial biotechnology platform for producing intrinsically toxic proteins and provides a foundation for their systematic biochemical, structural, and translational investigation.

## Results

### Standard recombinant expression methods are insufficient for highly toxic polymorphic toxins

Following the identification and functional validation of the PT families (Nachmias, et al., 2024), we sought to express PT1^Em^ (PT1 from *Escherichia marmotae,* hereafter PT1), PT7^Bc^ (PT7 from *Bacillus cereus*, hereafter PT7), PT8^Li^ (PT8 from *Leptospira interrogans*, hereafter PT8), and PT9^Cn^ (PT9 from *Cedecea neteri*, hereafter PT9) as recombinant proteins for biochemical and structural characterization. These PT families were selected because they belong to the subset of toxins for which cognate immunity proteins have been identified (Nachmias, et al., 2024), enabling evaluation of a toxin-immunity fusion strategy for recombinant production. Because these toxins target essential bacterial processes and exhibit potent antibacterial activity, their recombinant production posed substantial technical challenges due to host toxicity.

The first step in all the expression strategies described below involves molecular cloning of the toxin genes into plasmids compatible with the intended expression system. Since cloning and plasmid amplification occur in bacteria, even low-level basal toxin expression can be lethal to host cells, often resulting in plasmids carrying mutations within the toxin gene. To prevent this, all bacterial growth media used for molecular work contain 2% glucose to repress gene expression. Despite these precautions, basal promoter leakage remained sufficient to select for inactivating mutations or prevent stable propagation of several PT expression constructs, illustrating the exceptional toxicity of these proteins.

To evaluate recombinant expression, the genes encoding PT1, PT7, PT8, and PT9 were cloned as His-tagged constructs into the arabinose-inducible pBAD vector (see Materials and Methods). This vector exhibits relatively low basal expression and was previously used for toxicity validation assays of the PTs (Nachmias, et al., 2024). In parallel, the genes were cloned into IPTG-inducible pET28 vectors, which support high-level protein production but exhibit higher basal expression. Transformation of the pBAD constructs into E. coli BL21 cells yielded colony numbers comparable to those obtained with control plasmids. Transformation of the pET28 plasmids into BL21 cells resulted in either a complete absence (PT1) or a markedly reduced number of colonies compared with control plasmids, despite growth under repression conditions. Small-scale induction experiments followed by Western blot analysis failed to detect recombinant PT expression from either vector (Figure S1A).

Because co-expression of cognate immunity proteins has been successfully used to enable recombinant production of toxic bacterial effectors (Piscotta, et al., 2019, Hagan, et al., 2023), we next asked whether this strategy could overcome the expression barrier for PT proteins. We therefore applied this strategy to co-express PT1, PT7, PT8, and PT9 with their cognate immunity proteins (PIMs). To facilitate co-expression, toxin and immunity gene pairs were cloned into the dual-expression vector pETDuet (see Methods). Initial cloning and expression attempts were largely unsuccessful: PT7-PIM7 and PT9-PIM9 could not be stably cloned, suggesting persistent toxicity, while PT1-PIM1 and PT8-PIM8 were successfully cloned but showed no detectable expression (Figure 1A, B). Next, we evaluated a co-transformation strategy in which PT and PIM proteins were expressed from separate plasmids. PT8 and PT9 were cloned as His-tagged proteins into a pBAD24-p15Aori vector (modified pBAD24 with p15A origin, see Methods) to enable tightly regulated expression, while PIM8 and PIM9 were cloned as Strep-tagged proteins into the pET28 vector. Both plasmids were co-transformed into BL21 cells. To minimize toxin-mediated toxicity during induction, immunity protein expression was induced before toxin expression to enable intracellular neutralization of PT activity. Co-expression of PT8-PIM8 and PT9-PIM9 showed no detectable PT expression, whereas the PIM proteins were expressed (Figure 1B, C). These findings indicate that co-expression of cognate immunity proteins alone was insufficient to overcome the production barrier, suggesting that even transient accumulation of unbound toxin is sufficient to impair recombinant expression.

**Figure 1.**
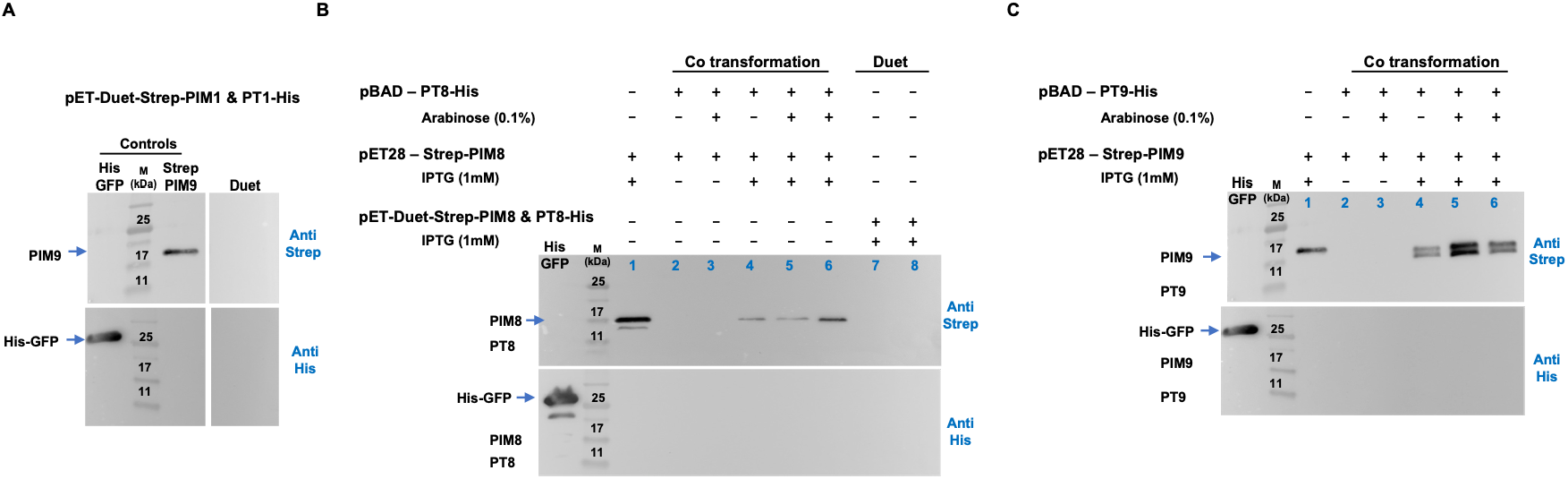
Expression of the PTs using standard recombinant expression approaches. **(A)** Co-expression of His-tagged PT1 and Strep-tagged PIM1 from the dual-expression vector pETDuet. Western blot analysis showed no detectable expression of PT1 or PIM1. **(B)** Co-expression of His-tagged PT8 and Strep-tagged PIM8 using either co-transformation of pBAD-PT8 and pET-PIM8 (lanes 2–6) or the pETDuet vector (lanes 7–8). In co-transformation experiments, PIM8 expression was induced either simultaneously with PT8 (lane 5) or 2 h before PT8 induction (lane 6). Western blot analysis showed no detectable PT8 expression under either condition. **(C)** Co-expression of His-tagged PT9 and Strep-tagged PIM9 using co-transformation of pBAD-PT9 and pET-PIM9. PIM9 expression was induced either simultaneously with PT9 (lane 5) or 2 h before PT9 induction (lane 6). Western blot analysis showed no detectable PT9 expression.

Finally, we asked whether eliminating host-cell viability constraints altogether could overcome the production barrier. Cell-free (CF) expression bypasses host-cell viability constraints, making it suitable for toxin expression (Ramm, et al., 2024, Woelbern and Ramm, 2024). PT1 could not be stably cloned into pIVEX plasmids for CF expression. In contrast, PT7, PT8, and PT9 were successfully cloned into pIVEX plasmids and expressed in the bacterial CF lysate. Western blot analysis showed no expression of the PTs (Figure S1B). These findings indicate that eliminating host-cell viability constraints alone is insufficient for the recombinant production of these toxins, suggesting that the newly synthesized proteins directly interfere with the bacterial cell-free transcription-translation machinery before detectable protein accumulates.

Collectively, these experiments demonstrate that neither reducing basal expression, co-expressing cognate immunity proteins, nor eliminating host-cell viability is sufficient to enable recombinant production of these highly toxic PT proteins. These findings motivated the development of an alternative production strategy that maintains toxin neutralization throughout the recombinant expression process.

### Fusion-based strategy for the production of wild-type PTs

To overcome production barriers associated with highly toxic PTs, we developed a fusion-based toxin neutralization strategy in which the PIM is covalently linked to the wild-type (wt) PT within a single polypeptide chain. We reasoned that covalently linking the toxin and immunity protein would promote rapid intramolecular complex formation during translation, thereby minimizing transient accumulation of unbound cytotoxic toxin. The fusion construct is organized as follows: an N-terminal Strep-tagged PIM is connected via a flexible linker to a C-terminal His-tagged PT (Figure 2A). The linker comprises multiple Gly-Gly-Gly-Ser (GGGS) repeats that provide conformational flexibility and facilitate formation of the predicted PIM-PT interaction interface (described below). Following the linker is a Tobacco Etch Virus protease cleavage site (TEVcs), which enables precise separation of the two domains after purification. The overall workflow was designed to maintain toxin neutralization throughout recombinant expression and initial purification while enabling subsequent recovery of the isolated PT protein following proteolytic cleavage and affinity-based purification. This strategy was intended to generate native-like PT proteins suitable for downstream biochemical, structural, and mechanistic analyses, as well as for future use as antimicrobials. TEV protease was selected because its highly specific cleavage generates the native PT with only a minimal N-terminal scar (one amino acid), thereby minimizing potential effects on protein function. (Figure 2A) (Carrington and Dougherty, 1988, Dougherty, et al., 1989, Polayes, et al., 1998). This minimal scar is particularly advantageous for downstream biochemical, structural, and mechanistic studies, where additional N-terminal residues can affect protein folding and activity.

**Figure 2.**
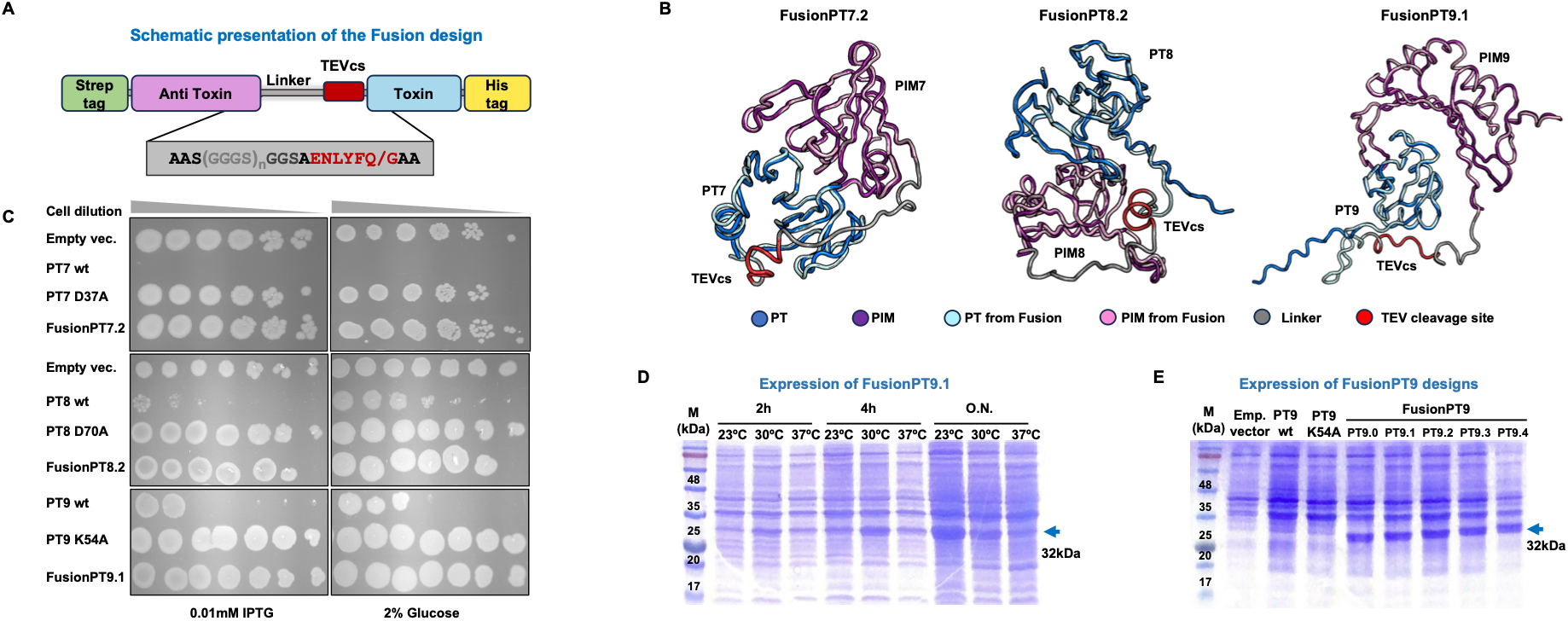
Structure-guided design of FusionPT constructs. **(A)** Schematic illustration of the fusion-based construct. The construct consists of an N-terminal Strep-tagged PIM linked to a C-terminal His-tagged PT through a flexible Gly-Gly-Gly-Ser (GGGS) linker containing a Tobacco Etch Virus protease cleavage site (TEVcs). (**B**) The selected FusionPT constructs for PT7, PT8, and PT9 expression. Predicted structures of the native non-covalent PT-PIM complexes were superimposed on those of the FusionPT.0-FusionPT.4 constructs containing GGGS linkers of varying lengths (Figure 2SB). The optimal fusion design was chosen as the shortest linker that preserved the relative orientation of the PT and PIM domains while minimizing structural distortion. FusionPT7.2, FusionPT8.2, and FusionPT9.1 were selected for experimental validation. **(C)** Fusion-based neutralization rescues PT toxicity. Drop assays of *E. coli* BL21 cells expressing FusionPT7.2, FusionPT8.2, and FusionPT9.1 under inducing conditions. Catalytically inactive PT mutants served as non-toxic controls. Fusion protein expression restored bacterial growth to levels comparable to those of the inactive mutants, indicating efficient neutralization of PT toxicity by the covalently linked PIM. **(D)** Optimization of fusion protein expression conditions. SDS-PAGE analysis of small-scale FusionPT9.1 expression at different induction temperatures and expression times. Overnight expression at lower temperatures produced the highest protein yield and was used for subsequent purification experiments. **(E)** Effect of linker length on FusionPT9 expression. SDS-PAGE analysis of FusionPT9 variants containing linkers of different lengths demonstrated comparable recombinant expression levels.

### Structure-guided design of PIM-PTwt fusion proteins

Because no experimental structures of the PT-PIM complex were available, linker design required computational prediction (e.g., Robetta, Boltz-2, and AlphaFold2 (Kim, et al., 2004, Jumper, et al., 2021, Passaro, et al., 2025)) of the relative orientation between the toxin and immunity protein. The linker must be sufficiently long to permit proper folding and intramolecular assembly of the PT-PIM complex, while remaining short enough to avoid introducing an excessively flexible, structurally heterogeneous region.

We used Robetta (Kim et al., 2004) to model the non-covalent PT-PIM complexes using deep learning-based structure prediction and generated a series of fusion constructs containing one to five GGGS repeats (FusionPT.0-FusionPT.1, Figure S2A). Each construct was compared with the predicted native complex, and the shortest linker that preserved the native domain arrangement was selected for experimental validation. Based on this workflow, we designed fusion variants of PT7, PT8, and PT9 (Figure 2B and Figure S2B). The same strategy was also applied to PT1. However, repeated cloning attempts predominantly yielded inactivating mutations within the PT1 coding sequence, consistent with the exceptional toxicity of this toxin even in the context of the fusion construct. Structural comparison of the predicted fusion constructs with their corresponding non-covalent PT-PIM complexes revealed that shorter linkers often imposed geometric constraints that distorted the relative orientation of the two domains (e.g., FusionPT9.0 and FusionPT7.0), whereas longer linkers provided no substantial improvement in structural agreement. FusionPT7.2, FusionPT8.2, and FusionPT9.1 were selected for experimental validation.

To evaluate whether the fusion designs alleviate PT-mediated toxicity during bacterial expression, we cloned the selected fusion constructs into the pET28 expression vector and transformed the resulting plasmids into E. coli BL21 cells. Toxicity was assessed by drop assay under inducing conditions. As controls, we included catalytically inactive PT mutants that have been shown to abolish toxicity and permit bacterial growth (Nachmias et al., 2024). Consistent with the proposed mechanism, expression of the fusion constructs restored bacterial viability to levels comparable to those of the non-toxic PT mutants, demonstrating that covalent linkage of the PIM effectively neutralizes PT toxicity during expression (Figure 2C).

To further evaluate the fusion design strategy, we generated a series of FusionPT9 constructs with linkers of varying lengths (FusionPT9.0-FusionPT9.4) and assessed toxicity using drop assays (ig. S2C). All fusion constructs restored bacterial growth. Notably, FusionPT9.1 consistently exhibited the strongest growth phenotype. These findings are consistent with the computational prediction that the shortest structurally compatible linker provides efficient intramolecular toxin neutralization.

### Expression of FusionPT7.2, FusionPT8.2, and FusionPT9.1

In our previous study, several catalytically inactive PT mutants lost their toxic phenotype yet failed to accumulate as recombinant proteins (Nachmias, et al., 2024). We therefore asked whether the fusion-based neutralization strategy not only alleviates toxicity but also enables productive recombinant expression. In contrast to all conventional expression strategies tested above (Figure 1), small-scale induction readily produced detectable levels of FusionPT7.2, FusionPT8.2, and FusionPT9.1 proteins (Figure S2D), demonstrating that intramolecular toxin neutralization enables the recombinant production of these highly toxic PTs. We next optimized expression conditions for large-scale production. Overnight induction produced substantially higher protein yields than shorter induction periods and was therefore used for all subsequent purification experiments (Figure 2D and Figure S2D). Comparison of the different FusionPT9 variants revealed no substantial differences in expression level, indicating that linker length primarily influences toxin neutralization (Figure 2C and Figure S2C) rather than recombinant protein accumulation under the conditions tested (Figure 2E).

These findings demonstrate that intramolecular toxin neutralization not only restores bacterial viability but also overcomes the previously insurmountable barrier to recombinant production of these highly toxic proteins.

### Purification of FusionPT7.2, FusionPT8.2, and FusionPT9.1

Having established robust expression of the fusion constructs, we next asked whether the fusion strategy could yield purified native toxins suitable for downstream biochemical and structural studies. To test this, the fusion proteins were purified, proteolytically cleaved to release the toxin domain, dissociated under denaturing conditions, and the isolated PTs were refolded and analyzed by size-exclusion chromatography (SEC). We first applied this workflow to FusionPT9.1. Strep-trap affinity purification yielded high levels of soluble fusion protein, and SDS-PAGE analysis indicated substantial enrichment of FusionPT9.1 after the affinity step (Figure 3A, B). Subsequent incubation with TEV protease resulted in a complete cleavage of the fusion construct, generating the expected PIM and PT products (Figure 3B). To dissociate the cleaved PIM-PT complex, the sample was incubated in 8 M urea and loaded onto a Ni^2+^-affinity column under denaturing conditions (Piscotta, et al., 2019, Hagan, et al., 2023). Because the PT domain retained its C-terminal His-tag, it could be selectively captured on the resin, while the dissociated PIM and other contaminants were removed during washing. Gradual urea removal directly on the column enabled refolding of the immobilized PT protein prior to elution (Figure 3C). Finally, the refolded protein was subjected to SEC, which revealed a predominantly monodisperse monomeric peak consistent with the successful recovery of soluble, properly folded PT9wt (Figure 3D). This procedure yielded highly purified PT9wt, as indicated by SDS-PAGE analysis (Figure 3E).

**Figure 3.**
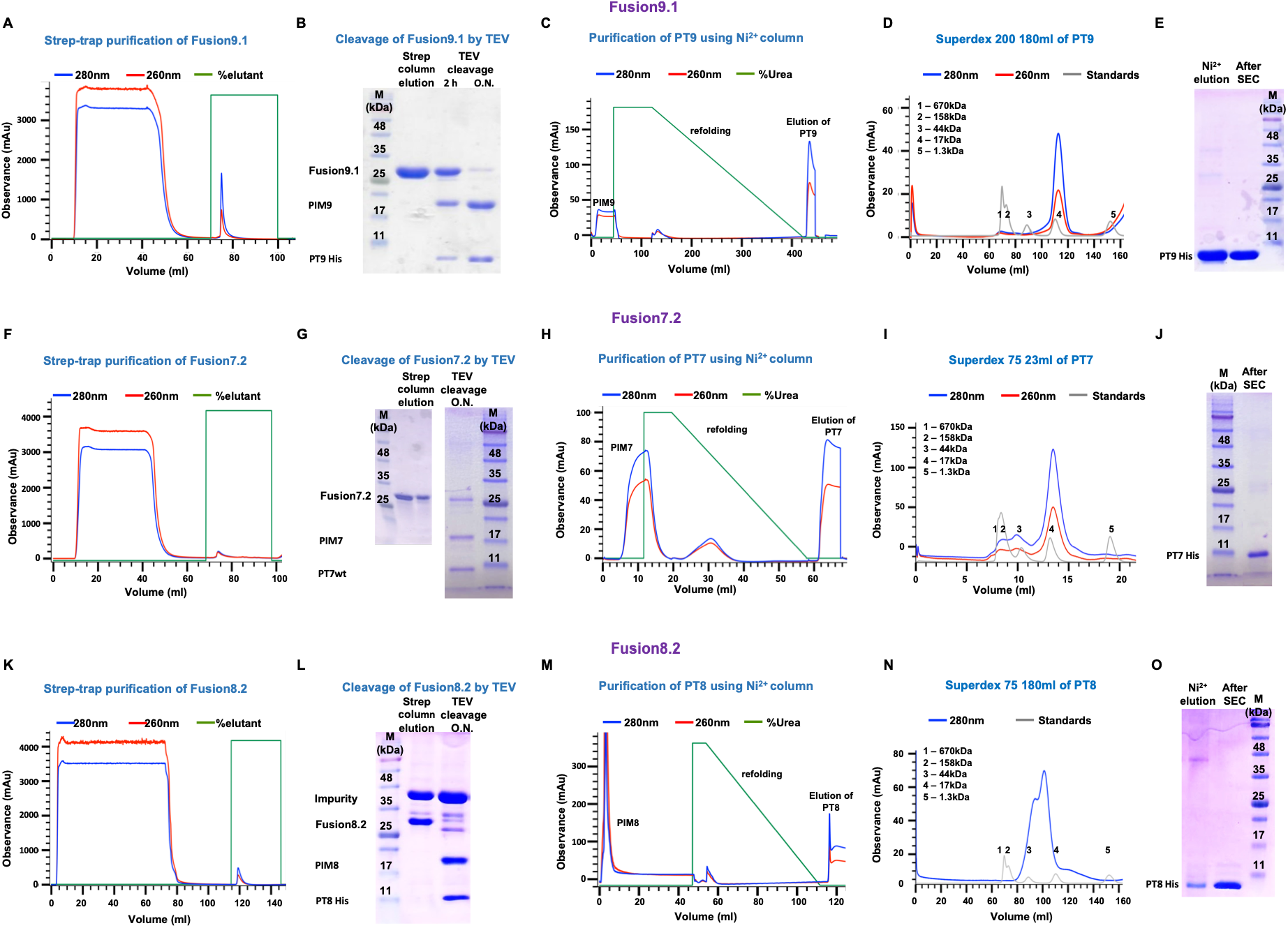
Purification of wild-type PT7, PT8, and PT9 using the fusion-based expression strategy. Purification of PT9 **(A-E)**, PT7 **(F-J),** and PT8 **(K-O).** (**A, F, K**) Strep-tag affinity purification of FusionPT9.1, FusionPT7.2, and FusionPT8.2. (**B, G, L**) In vitro TEV protease cleavage of the purified Fusion protein. Cleavage was assessed by SDS-PAGE. FusionPT8.2 was co-purifying with a ∼50-kDa contaminant; the contaminating protein remained uncleaved (**L**). (**C, H, M**) Purification of the wt PTs by Ni^2+^-affinity chromatography following denaturation of the cleaved PIM-PT complex in 8 M urea, on-column refolding, and elution under native conditions. (**D, I, N**) SEC profile of purified PTs. (**E, J, O**) SDS–PAGE analysis of the purified PT9 following SEC indicating high purity.

Next, we applied the same workflow to FusionPT7.2. Strep-tag affinity purification yielded substantial amounts of FusionPT7.2 (Figure 3F-J). TEV protease treatment resulted in near-complete cleavage of the fusion construct (Figure 3G). Subsequent dissociation of the PIM-PT complex under denaturing conditions, followed by Ni^2+^-affinity purification and SEC, enabled recovery of highly purified PT7 (Figure 3H-J). Similarly, Strep-tag affinity purification yielded substantial amounts of FusionPT8.2 protein (Figure 3K-L). Unlike PT7 and PT9, purification of FusionPT8.2 co-purified an additional ∼50-kDa protein (Figure 3K-L). TEV protease treatment of the eluted proteins resulted in complete cleavage of FusionPT8.2, while the co-purifying protein remained unaffected, consistent with its non-specific origin (Figure 3L). Following dissociation of the PIM-PT complex under denaturing conditions and Ni^2+^-affinity purification, PT8 toxin was efficiently separated from the contaminating protein and recovered at high purity (Figure 3M-O).

Together, these results demonstrate that the fusion-based purification strategy is robust across multiple polymorphic toxin families and enables recovery of highly purified wt toxins despite differences in purification behavior and cleavage efficiency. Importantly, the strategy consistently yielded soluble proteins that could be successfully refolded after denaturation, providing material suitable for downstream biochemical and structural analyses.

### Intracellular Proteolysis Simplifies and Improves the Fusion Production Platform

Having established that the fusion-based strategy enables purification of multiple wt PT proteins, we next sought to simplify the workflow to improve scalability and reduce production costs. Although effective, the original purification protocol consisted of four steps performed over three days, including Strep-tag affinity purification and in vitro TEV protease cleavage, both of which substantially increase production costs and processing time (Figure 4A). To address these limitations, we developed an optimized purification strategy based on intracellular TEV-mediated cleavage of the fusion construct. We reasoned that the high affinity of the PT-PIM interaction would maintain the complex after intracellular TEV cleavage, thereby preserving toxin neutralization while eliminating the need to purify the intact fusion protein and to perform subsequent in vitro proteolysis. This strategy reduced the workflow to two purification steps and eliminated the need for both Strep-tag affinity purification and large-scale TEV protease treatment (Figure 4A).

**Figure 4.**
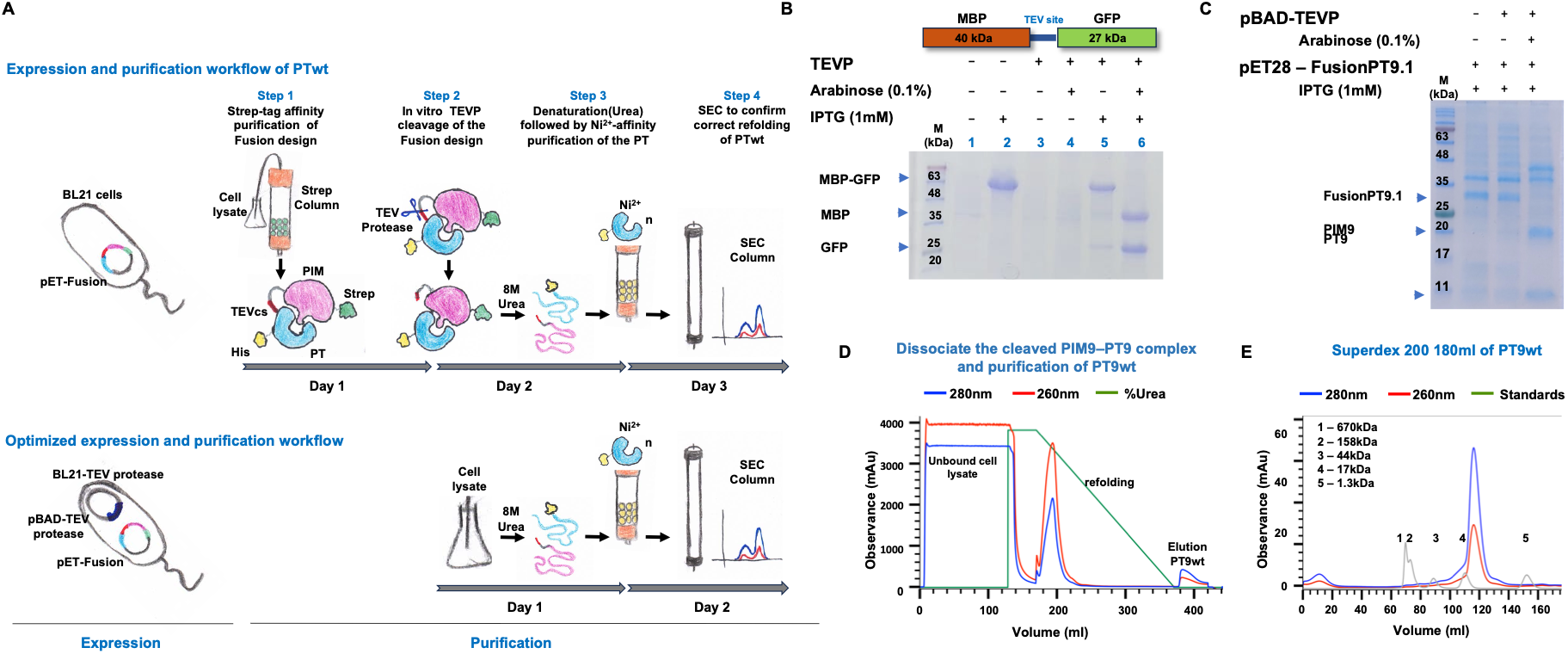
Optimization of the fusion-based purification workflow through intracellular TEV-mediated cleavage. **(A)** Schematic comparison of the original four-step purification workflow and the optimized two-step workflow. In the optimized protocol, intracellular TEV protease cleavage replaces both Strep-tag affinity purification and in vitro TEV cleavage, enabling direct purification of the released PT by Ni^2+^ affinity chromatography, followed by SEC. **(B)** Validation of intracellular TEV protease activity using the MBP-TEVcs-GFP reporter. BL21 cells or tBL21-TEVP cells were induced with IPTG alone or with IPTG and arabinose to co-induce expression of MBP-TEVcs-GFP and TEV protease. SDS-PAGE analysis demonstrates complete intracellular cleavage of the reporter protein upon TEV induction. **(C)** Intracellular cleavage of FusionPT9.1. FusionPT9.1 was co-expressed with TEV protease in tBL21-TEVP cells. SDS-PAGE analysis demonstrates efficient cleavage of the fusion construct into the expected products. **(D)** Purification of PT9 using the optimized workflow. Following intracellular TEV-mediated cleavage, soluble lysates were denatured in 8 M urea, and PT9 was purified by Ni^2+^-affinity chromatography with on-column refolding. **(E)** SEC profile of purified PT9 obtained using the optimized purification workflow, showing recovery of a predominantly monodisperse protein consistent with successful refolding. SDS-PAGE analysis confirms the high purity of PT9.

To enable intracellular cleavage, we engineered an arabinose-inducible TEV protease expression plasmid compatible with the pET expression system by replacing the pBR322 origin of pBAD24 with the p15A origin (Figure S3A). The resulting pBAD-TEVP plasmid was introduced into BL21cells to generate the tBL21-TEVP strain used throughout this study (Figure S3A).

To evaluate in-cell proteolytic activity, tBL21-TEVP competent cells were transformed with the pET28-MBP-TEVcs-GFP plasmids. As a control, the plasmid was transformed into BL21 cells. Induction of MBP-GFP led to robust accumulation of the fusion protein in both strains (Figure 4B, wells 2 and 5). In contrast, co-induction of TEV protease in tBL21-TEVP cells resulted in complete cleavage of the MBP-GFP fusion protein (Figure 4B, well 6). We next examined whether TEVP could efficiently cleave the FusionPT9.1 construct in vivo. To this end, pET28-FusionPT9.1 was transformed into tBL21-TEVP cells, and expression of both FusionPT9.1 and TEVP was induced simultaneously. SDS-PAGE analysis revealed efficient cleavage of the fusion construct, yielding the expected cleavage products and demonstrating that intracellular TEV proteolysis is compatible with the FusionPT9 architecture (Figure 4C).

Having demonstrated efficient intracellular cleavage of FusionPT9.1, we next asked whether this simplified workflow could recover purified native PT9. FusionPT9.1 was co-expressed with TEVP in tBL21-TEVP cells, and the cells were then harvested and lysed. The soluble fraction was isolated and incubated with 8 M urea to dissociate the cleaved PIM-PT complex. The denatured sample was loaded directly onto a Ni2+-affinity column and gradually washed with buffer to promote refolding of the immobilized protein before elution (Figure 4D). SDS-PAGE analysis showed high purity of PT9 (Figure S3B). Finally, SEC analysis revealed a predominantly monodisperse peak, consistent with successful refolding and recovery of soluble PT9 (Figure 4E). Importantly, in addition to reducing the purification workflow from four steps to two, the optimized protocol increased PT9 recovery from approximately 1-2 mg/L to ∼4 mg/L of bacterial culture, representing a two- to four-fold increase in protein yield.

Together, these results demonstrate that intracellular TEV-mediated cleavage substantially simplifies the production platform without compromising purification efficiency. In addition to eliminating two purification steps and reducing reliance on affinity reagents and exogenous protease, the optimized workflow increased recombinant protein recovery, making the platform more suitable for scalable production of highly toxic bacterial toxins.

### Computational redesign of immunity proteins expands the fusion-based production platform

Although the fusion-based strategy enabled efficient production of multiple wt PT proteins, several PIM-PT fusion constructs still failed to yield detectable recombinant protein. We hypothesized that, for these proteins, the native PIM does not bind its cognate toxin with sufficient affinity to fully suppress toxin activity during expression, allowing residual toxicity to impair recombinant protein production. As a proof of concept, we selected PT37, a previously uncharacterized polymorphic toxin from *E. coli* that consistently failed recombinant production. To overcome this limitation, we sought to engineer improved toxin-binding proteins through computational protein design. The approach was based on the structural features of the naturally occurring immunity proteins and sought to improve their interaction with the toxin domain. Consistent with other highly toxic PTs, recombinant expression of PT37 yielded no detectable protein (Figure S4A), despite only a relatively mild toxic phenotype in drop assays (Figure 5A).

**Figure 5.**
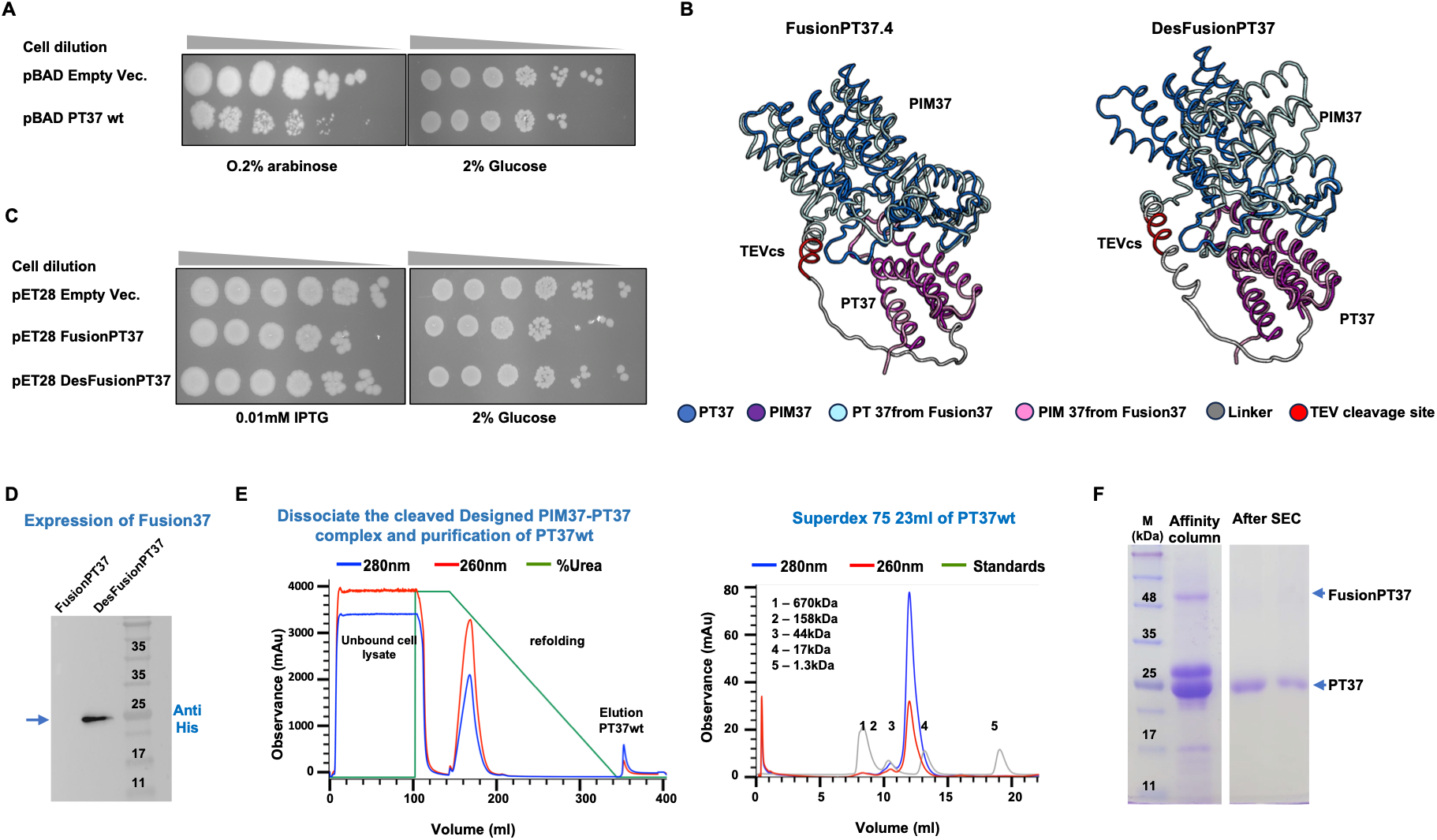
Computational redesign of immunity proteins expands the fusion-based production platform. **(A)** Drop assay demonstrating the toxicity of PT37. **(B)** Structure-guided design of the FusionPT37 and DesFusionPT37. Predicted structure of the native non-covalent PT37-PIM37 complex was superimposed with the corresponding FusionPT37 and DesFusionPT37 predicted structures. **(C)** Drop assay showing rescue of PT37 toxicity by FusionPT37 and DesFusionPT37. **(D)** Western blot analysis of small-scale expression of FusionPT37 and DesFusionPT37 indicating recombinant expression of DesFusionPT37 but not FusionPT37. **(E)** Purification of PT37 using the optimized intracellular TEV-mediated purification workflow. **(H)** SDS-PAGE analysis of purified PT37.

We first applied our fusion-based strategy by generating a FusionPT37 construct in which PT37 was fused to its predicted cognate immunity protein identified using AlphaFold2 (Jumper, et al., 2021) (Figure 5B, left). Although the fusion construct efficiently rescued the toxic phenotype in drop assays (Figure 5C), no recombinant protein was detected in small-scale expression experiments (Figure 5D, left), indicating that the native immunity protein was unable to fully neutralize PT37 during expression. We therefore sought to improve toxin neutralization by engineering the immunity protein rather than modifying the toxin itself. Starting from the predicted PT37-PIM37 complex, ProteinMPNN (Dauparas et al., 2022) was used to redesign the immunity protein and linker while preserving the overall structure of the immunity protein (Figure S4B) and the complex (Figure 5B), with the goal of strengthening the toxin-immunity interaction and promoting productive recombinant expression. (Figure S4C)

The resulting redesigned fusion construct, DesFusionPT37, was subsequently evaluated experimentally. Drop assays demonstrated efficient rescue of PT37-mediated toxicity, while small-scale expression experiments revealed production of the recombinant protein, which was not observed with the original FusionPT37 construct (Figure C, D). Next, DesFusionPT37 was expressed in tBL21-TEVP cells and purified using the optimized purification protocol (Figure 5E). SDS-PAGE analysis demonstrated high purity of PT37 (Figure 5F). These findings demonstrate that computational redesign of natural immunity proteins can rescue recombinant production when native toxin-immunity pairs are insufficient. Thus, computational engineering substantially expands the applicability of the fusion-based production platform beyond naturally optimized toxin-immunity systems.

### Functional Validation of Purified PTwt Proteins

To determine whether the fusion-based production strategy yielded correctly folded and enzymatically active proteins, we examined the nuclease activity of the purified PTs. We have previously demonstrated nuclease activity of PT7 using PT7 mutants that abolish in vivo toxicity but retain enzymatic activity in vitro (Nachmias, et al., 2024). We therefore asked whether PTwt proteins produced using our workflow retained their enzymatic activity despite being purified as PT-PIM complexes and then denatured and refolded.

Purified PT7, PT8, and PT9 were incubated with plasmid DNA and DNA degradation was analyzed by agarose gel electrophoresis. DNase A served as a positive control. Consistent with our previous findings, purified PT7 exhibited robust nuclease activity, resulting in extensive degradation of the DNA substrate (Figure 6A). In addition to confirming the previously reported DNase activity of PT7, these experiments identified PT8 as a second nuclease within this family. In contrast, no detectable nuclease activity was observed for PT9 (Figure 6A).

**Figure 6.**
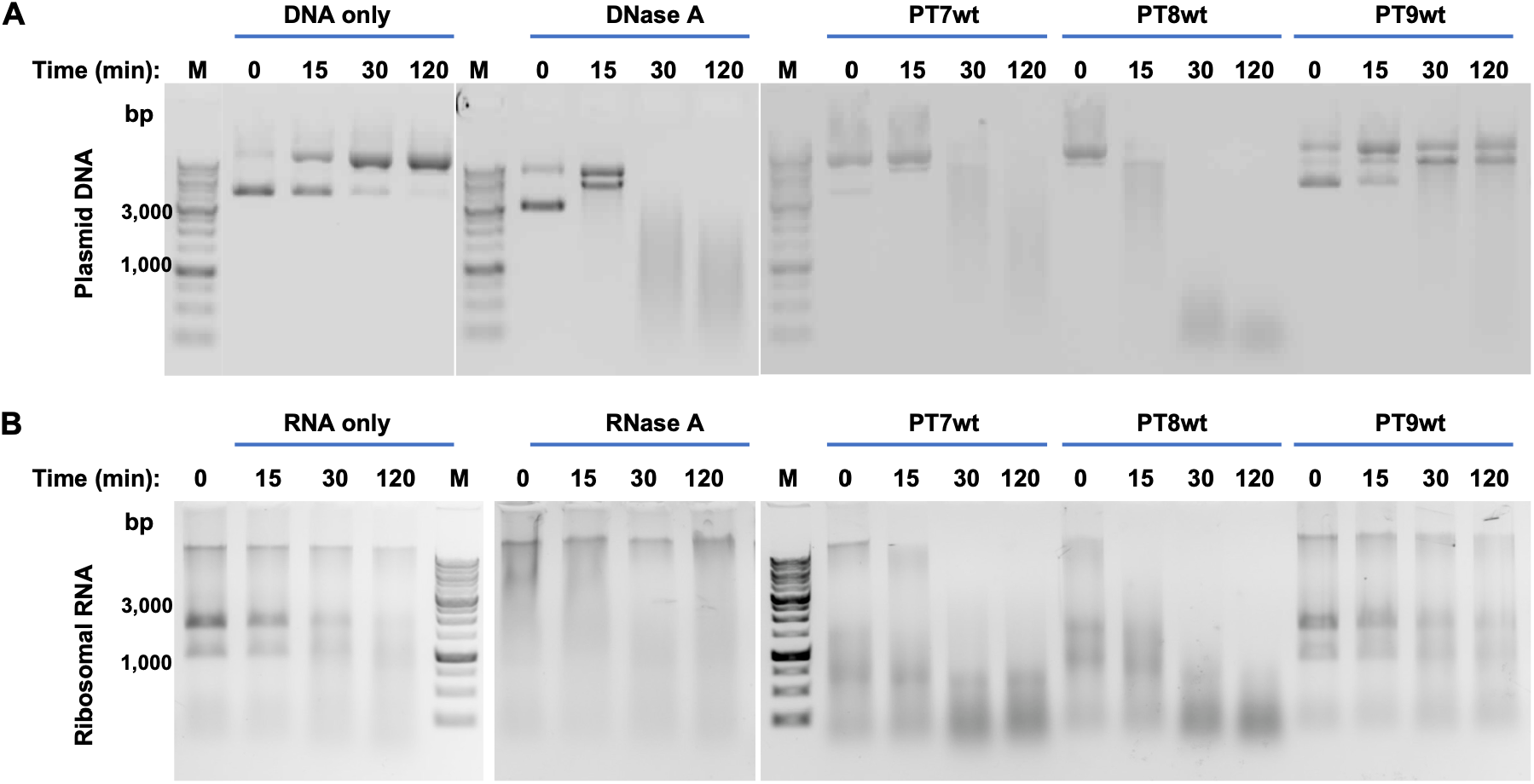
Functional validation of purified wild-type PT proteins. **(A)** In vitro nuclease activity assay of purified PT7, PT8, and PT9. Purified plasmid DNA was incubated with the indicated proteins for 0-120 min at 37 °C, and DNA degradation was analyzed by 1% agarose gel electrophoresis. Buffer alone and DNase A served as the negative and positive controls, respectively. **(B)** In vitro RNase activity assay of purified PT7, PT8, and PT9. Purified ribosomal RNA was incubated with the indicated proteins under the same reaction conditions as in (A), and RNA degradation was analyzed by agarose gel electrophoresis.

To further investigate the substrate specificity of the purified toxins, we next examined their ability to degrade ribosomal RNA. We previously demonstrated that the catalytically active PT7 mutant does not degrade tRNA (Nachmias, et al., 2024), indicating that its RNase activity is substrate selective rather than indiscriminate. We therefore asked whether PT7, PT8, and PT9 target ribosomal RNA. Purified PT7, PT8, and PT9 were incubated with purified bacterial ribosomal RNA (rRNA) under identical reaction conditions, and RNA integrity was analyzed by agarose gel electrophoresis. Both PT7 and PT8 efficiently degraded ribosomal RNA, whereas PT9 showed no detectable RNase activity (Figure 6B). These findings indicate that PT7 and PT8 possess both DNase and rRNase activities under the conditions tested, whereas PT9 showed no detectable nuclease activity toward either substrate.

Together, these findings demonstrate that the fusion-based production platform preserves the native enzymatic activity of highly toxic proteins, even after transient toxin neutralization, denaturation, and refolding during purification. The successful recovery of catalytically active PT7 and PT8 validates the platform as a reliable approach for producing functional bacterial toxins suitable for biochemical, structural, and mechanistic studies. Furthermore, identifying PT8 as a nuclease illustrates how this platform enables the functional characterization of previously inaccessible toxin families.

## Discussion

The rapid emergence of AMR has intensified the search for antimicrobial agents with novel mechanisms of action (Antimicrobial Resistance, 2022). The recombinant production of intrinsically toxic proteins has long represented a major challenge in protein engineering, structural biology, and antimicrobial discovery. Although bacterial protein toxins are among the most biologically and therapeutically important examples, their inability to be produced has severely limited mechanistic investigation and translational development. In this study, we establish a modular and scalable production platform that overcomes this bottleneck through transient intramolecular neutralization during recombinant expression. By integrating toxin-immunity fusion, computational protein engineering, and streamlined purification, we successfully produced multiple previously intractable polymorphic toxins while preserving their native enzymatic activity.

Multiple conventional recombinant expression strategies failed to produce detectable amounts of PT proteins (Figure 1). Neither tightly regulated inducible promoters, co-expression of cognate immunity proteins, nor bacterial cell-free expression systems enabled productive recombinant expression. These findings emphasize that the primary obstacle is not simply host viability but rather the rapid activity of newly synthesized toxin molecules during protein production. Even transient accumulation of a small fraction of unneutralized toxin appears sufficient to prevent protein accumulation.

An interesting question raised by these findings is why recombinant co-expression of toxin-immunity pairs fails, even though these proteins are naturally co-produced in their native bacterial hosts. Several non-mutually exclusive explanations may account for this discrepancy. First, recombinant expression from multicopy plasmids under inducible promoters is likely to generate toxin molecules at rates that substantially exceed physiological levels, increasing the probability that toxin molecules transiently escape neutralization before immunity proteins can bind. Second, several bacterial toxins exhibit catalytic activity for which even a single active molecule may be sufficient to irreversibly inhibit essential cellular processes, thereby preventing continued protein synthesis and cell growth (Johnson, et al., 2013). Third, polymorphic toxins are typically synthesized via specialized secretory pathways and are frequently associated with dedicated secretion machinery, trafficking factors, or cellular scaffolds prior to export(Halvorsen, et al., 2024). These native regulatory mechanisms are absent during heterologous expression in *E. coli*, potentially exposing the host cytoplasm to toxin molecules in a manner that does not occur under physiological conditions.

Our observation that bacterial cell-free expression also failed to produce detectable toxin provides additional mechanistic insight into the production barrier (Figure S1B). Cell-free protein synthesis has been widely used for recombinant production of toxic proteins because it eliminates host-cell viability constraints while retaining the bacterial transcription-translation machinery (Woelbern and Ramm, 2024). The failure of PT expression under these conditions therefore suggests that cellular toxicity alone is not responsible for the production defect. Instead, newly synthesized toxin molecules may directly inhibit the transcription-translation machinery present within the lysate. Because PT7 and PT8 exhibit DNase and RNase activities (Figure 6), they may rapidly degrade ribosomal RNA, messenger RNA, or other nucleic acid components required for continued protein synthesis, thereby terminating translation before detectable protein accumulates. Although this mechanism remains to be experimentally validated, our findings suggest that eliminating host viability alone is insufficient for recombinant production of highly toxic enzymatic proteins. Rather, toxin activity must remain effectively neutralized throughout protein synthesis.

The conceptual advance of this work lies in converting a naturally occurring intermolecular regulatory interaction into a broadly applicable strategy for recombinant protein production. Rather than attempting to suppress toxin expression or tolerate its activity, our platform transiently neutralizes toxicity during biosynthesis while allowing complete recovery of the native protein following purification. Because this strategy is independent of the toxin’s molecular target or catalytic mechanism, we anticipate that it will be broadly applicable to diverse classes of intrinsically toxic proteins.

An important aspect of the platform is its integration with modern computational protein modeling. Because none of the PT-PIM systems examined here had experimentally determined structures, linker design relied entirely on deep learning-based structure prediction. The agreement between computational predictions and experimental outcomes illustrates how recent advances in protein structure prediction can substantially accelerate construct optimization for previously uncharacterized protein families. More broadly, this study demonstrates how computational structural biology can be integrated directly into recombinant protein engineering workflows rather than being used solely for structural interpretation.

A second important advance is the incorporation of computational protein design into the production platform. Although fusing to the native immunity protein enabled production of several PT families, this strategy was insufficient for PT1 and PT37, indicating that naturally evolved immunity proteins are not always optimal for recombinant overexpression. Importantly, redesign of the PT37 immunity protein using ProteinMPNN (Figure 5) converted a nonproductive construct into one that supported robust recombinant expression and purification. This finding transforms the platform from one that simply exploits naturally occurring toxin-immunity interactions into one that can actively engineer them. As computational protein-design methods continue to improve, including approaches such as ProteinMPNN (Dauparas et al., 2022), PROSS (Weinstein, et al., 2021), RFdiffusion (Watson, et al., 2023), and related algorithms, we anticipate that affinity, stability, and specificity of toxin-binding proteins can be further optimized, extending the applicability of this strategy to increasingly challenging toxin families. The inability to express PT1 even using the fusion strategy likely reflects an extreme case in which the native toxin-immunity interaction remains insufficient to fully suppress toxicity under recombinant overexpression conditions. Nevertheless, the successful rescue of PT37 demonstrates that such limitations are not necessarily intrinsic to the platform but can potentially be overcome through rational engineering of toxin-binding partners.

Beyond improving expression, we also substantially simplified the purification workflow. Intracellular TEV-mediated cleavage eliminated two purification steps while increasing protein yield approximately two- to four-fold. This reduction in processing time, reagent consumption, and production costs substantially improves the platform’s scalability and practical utility for recombinant protein production, facilitating larger-scale manufacturing for structural biology, biochemical characterization, high-throughput screening, and industrial biotechnology.

More broadly, we propose transient intramolecular neutralization as a general strategy for the recombinant production of intrinsically toxic proteins. Although demonstrated here with polymorphic toxins, the platform is conceptually applicable to a much broader range of intrinsically toxic proteins whose biological activity interferes with recombinant production. Future integration with advances in computational protein design may further expand this strategy beyond naturally occurring toxin-immunity systems by enabling the engineering of synthetic neutralizing binders for proteins lacking cognate immunity partners. This work provides a scalable microbial biotechnology platform that should facilitate the recombinant production of previously inaccessible proteins for structural biology, synthetic biology, enzyme engineering, and antimicrobial discovery.

## Materials and Methods

### Plasmids Construction

The plasmids used in this project are as follows: the pET28a(+) vector (Novagen) was used for IPTG-inducible expression. This vector contains a T7 promoter for high-level expression and a pBR322 origin of replication (Ori). The pBAD24 vector (Thermo Fisher Scientific catalog number V43001) was used for tightly regulated arabinose-inducible expression. The vector contains the araBAD promoter, which enables low basal expression and tightly controlled induction conditions, and a pBR322 Ori. pBAD24-p15Aori, is a modified pBAD24 in which the pBR322 Ori was replaced with the p15A origin (Figure S3A). The pETDuet-1 vector (Novagen) was used for co-expression experiments involving toxin-immunity protein pairs. This dual-expression vector contains two multiple cloning sites, each driven by independent T7 promoters and ribosome-binding sites, enabling simultaneous expression of two recombinant proteins. The pIVEX2.4d vector was used for bacterial cell-free protein expression experiments. This vector is optimized for coupled transcription-translation reactions in E. coli-based CF lysates and contains a T7 promoter for in vitro protein synthesis.

Plasmids were constructed using the Gibson Assembly method (Gibson, et al., 2009). Vector backbones were linearized by PCR using primers flanking the multiple cloning site (Sigma-Aldrich, Israel). PT and PIM genes were amplified from plasmids generated in our previous study (Nachmias, et al., 2024), incorporating either a C-terminal 6×His tag (PTs) or an N-terminal Strep tag (PIMs). Genes encoding the FusionPT constructs were synthesized commercially (Twist Bioscience). The pET28-MBP-GFP plasmid was constructed by inserting the MBP-TEV site-GFP sequence with a C-terminal 6XHis Tag into the multiple cloning site of the pET28 plasmid.

DNA fragments were designed with 15–20 bp overlapping regions and assembled using Gibson Assembly. Reactions were performed in a total volume of 4 µL containing 50 ng of linearized vector and an equimolar amount of insert DNA. Assembly mixtures were incubated at 50°C for 1 h and subsequently transformed into chemically competent E. coli TOP10 cells by heat shock. Transformants were selected on LB agar supplemented with 2% glucose and the appropriate antibiotic, kanamycin for pET28-derived plasmids and ampicillin for pBAD-, pETDuet-1-, and pIVEX2.4d-derived plasmids. Individual colonies were screened and verified by Sanger sequencing.

### Construction of the pBAD-MBP-TEVP-p15A vector and generation of tBL21-TEVP competent cells

A synthetic DNA fragment encoding maltose-binding protein (MBP) followed by a TEV cleavage site and fused to tobacco etch virus protease (TEVP) was inserted into the multiple cloning site of pBAD-p15A (pBAD-MBP-TEVP-p15A vector, hereafter pBAD-TEVP) (Figure S3A). To generate transient TEVP-expressing BL21(DE3) competent cells (tBL21-TEVP), chemically competent BL21(DE3) cells were transformed with pBAD-TEVP and selected on LB agar supplemented with ampicillin. Individual transformants were used to prepare chemically competent tBL21-TEVP cells.

### Small-Scale Expression of the FusionPT proteins

Chemically competent BL21(DE3) cells were transformed with the target plasmid and selected on LB agar supplemented with the appropriate antibiotic. For small-scale expression, overnight culture was grown in LB medium supplemented with 2% glucose and the appropriate antibiotic. The next day, the culture was diluted 1:100 into fresh medium supplemented with the appropriate antibiotic and 0.1% glucose and incubated at 37 °C until OD₆₀₀ ≈ 0.5-0.6, and the inducer (IPTG, 1 mM or arabinose, 0.2%) was added. After overnight incubation at 23, 30, or 37 °C, 0.5 ml samples were harvested by centrifugation, lysed with BugBuster® Master Mix (Millipore, Cat. No. 71456) supplemented with 1 mM PMSF (Sigma-Aldrich, Israel) and Protease Inhibitor Cocktail Set V (Millipore, Cat. No. 539137). Soluble and insoluble fractions were separated by centrifugation and analyzed by SDS-PAGE.

### Protein Expression and Purification

For large-scale protein expression, cultures were grown and induced as described above for small-scale expression. Cells were harvested by centrifugation and resuspended in lysis buffer consisting of 20 mM Tris-HCl (pH 7.5), 100 mM NaCl, 5 mM MgCl₂, and 0.2% Triton X-100 supplemented with lysozyme and protease inhibitors. Cells were disrupted using a Cell Disruptor (Microfluidics LV1-1) and insoluble material was removed by centrifugation for 30 min at 4°C.

For the standard fusion protein purification workflow, clarified soluble lysates were loaded onto a 3-ml Strep-Tactin Sepharose column (Cytiva) pre-equilibrated with buffer containing 20 mM Tris-HCl (pH 7.5) and 100 mM NaCl (Tris buffer) using an ÄKTA chromatography system. Following washing, bound proteins were eluted with phosphate-buffered saline (PBS) supplemented with 2.5 mM desthiobiotin (Cayman chemical, 30418-500). Fusion proteins were subsequently incubated overnight at 4°C with recombinant TEV protease at an enzyme-to-substrate molar ratio of approximately 1:20 to cleave the linker separating the PIM and PT domains. The next day, TEV cleavage was verified by SDS-PAGE. Cleaved samples were denatured in 8 M urea for 3 h at room temperature and applied to a 5-mL HisTrap FF column (Cytiva). The column was washed with denaturing buffer to remove contaminants. Refolding of immobilized PT proteins was achieved by gradually replacing the denaturing buffer with Tris buffer directly on the column. Proteins were eluted with the Tris buffer supplemented with 250 mM imidazole. Eluted proteins were concentrated and further purified by SEC using a Superdex 75 Increase 10/300 GL column (Cytiva) equilibrated with Tris buffer. Fractions corresponding to the expected elution volume were pooled, concentrated, flash-frozen, and stored at −80°C.

### Optimized Purification Workflow Based on Intracellular Proteolysis

For the optimized purification workflow, intracellular TEV-mediated cleavage was performed during protein expression, eliminating the need for Strep-tag affinity purification and in vitro protease treatment. For protein expression, the fusion-containing plasmid was transformed into chemically competent tBL21-TEVP cells. Protein expression was performed as described above, except that IPTG and arabinose were added simultaneously to induce expression of both the fusion protein and TEV protease. Following cell lysis, soluble fractions were incubated in 8 M urea for 3 h and directly applied to a 5-mL HisTrap FF column (Cytiva). Bound proteins were washed under denaturing conditions and subsequently refolded on-column by gradual exchange into the Tris buffer. Proteins were eluted with Tris buffer supplemented with 250 mM imidazole, concentrated, and further purified by SEC using a Superdex 75 Increase 10/300 GL column (Cytiva). Fractions corresponding to monomeric protein were pooled, concentrated, aliquoted, and stored at −80°C.

### Protein Detection by Western Blot

Proteins were separated by SDS–PAGE and transferred onto nitrocellulose membranes. Membranes were blocked, washed, and incubated with the primary antibody, mouse anti-His (LifeTein, LLC LT0426), followed by incubation with HRP-conjugated goat anti-mouse secondary antibody (Jackson, 115-035-062). Detection of Strep-tagged proteins was performed using Precision Protein StrepTactin-HRP Conjugate (Bio-Rad, #1610381) according to the manufacturer’s instructions. Next, membranes were washed and developed using enhanced chemiluminescence (ECL; azure biosystems, AC2101). Chemiluminescent signals were detected and recorded using a ChemiDoc imaging system (Bio-Rad).

### Nuclease and RNase assay

To measure nuclease activity, 0.5 μg plasmid DNA (purified pAdVAntage plasmid; Promega), was incubated with the PT7, PT8, PT9, DNase A, or water (as a control) in reaction buffer containing 10 mM MgCl_2_, 100 μg/ml bovine serum albumin, and 10 mM Bis-Tris propane (pH 7) at 37 °C for different time periods. Nuclease reactions were analyzed on a 1.0% agarose gel and stained with ethidium bromide. RNase activity was measured under similar conditions using 0.5 μg of total ribosomal RNA extracted from mammalian cells.

### Drop Assay

Overnight culture was grown in LB medium supplemented with 2% glucose and the appropriate antibiotic. The next day, the culture was diluted 1:100 into fresh medium supplemented with the appropriate antibiotic and 0.1% glucose, incubated at 37 °C until OD_600_≈ 0.5-0.6. Cultures were then serially diluted tenfold (10⁰–10⁻⁸) in sterile 96-well plates, and 2 µL of each dilution was spotted in parallel onto repression plates (2% glucose) and induction plates (0.2% arabinose for pBAD or 0.01 mM IPTG for pET28a). Plates were incubated overnight at 37 °C, and growth was documented the following day. Drop assays were repeated in three independent biological experiments with comparable results. Representative images are shown.

### Cell free

#### Cell-free protein production

The proteins were produced in vitro in continuous exchange mode (CECF), with a reaction-to-dialysis volume ratio of 1:10. All reactions were performed at 27°C with gentle rotation at 18 rpm in a hybridisation oven (Hybrynogene K38, Ascon Technologic, Techne) using a 3 mL dialysis system (Maxi GeBaFlex, 8 kDa MWCO, Gene Bio-Application Ltd) placed in 50 ml Eppendorf tubes containing 30 mL of feeding solution. The reaction mixtures contained the following: target protein pIVEX plasmid (16µg/ml); *E. coli* BL21(DE3) S30 cellular extract; and T7 polymerase. Other standard cell-free components in the reaction and dialysis solutions (e.g. amino acids, NTPs, tRNAs, HEPES, MgOAc_2_, creatine phosphate, creatine kinase, NH_4_OAc, spermidine, cAMP, folinic acid) were added as described in a previously published protocol (Imbert et al 2021). The cell-free reaction was stopped after 16 hours. The reaction mixtures were then centrifuged for 20 minutes at 14,000 g prior to western blot analysis.

## Acknowledgements

This work is funded in part by the Israel Innovation Authority, grants #81260 (to N.T.) and #81259 (to A.L.). This work used the Cell-Free facility (Lionel Imbert) at the Grenoble Instruct-ERIC Center (ISBG; UAR 3518 CNRS-CEA-UGA-EMBL) within the Grenoble Partnership for Structural Biology (PSB). Platform access was supported by Instruct-ERIC PID: 25173.

## Author contributions

I.C., R.F., and N.T. conceived and designed the study; T.F., R.F., I.C., and N.D. performed the cloning and recombinant expression of wt PT proteins; N.T., I.C., R.F., and N.D. designed the FusionPT constructs; R.K. and S.B. performed the drop assays and small-scale expression experiments of the FusionPT constructs; I.C., R.F., T.S., and S.B. performed large-scale expression and purification of the FusionPT proteins; R.F. developed the bacterial TEV protease expression system; S.C., M.S., and A.L. discovered PT37 and validated its toxicity; T.S. designed the FusionPT37 construct; H.N. performed the nuclease assays; R.F., I.C., and N.T. analyzed the data and wrote the manuscript with input from all authors. All authors reviewed and approved the final manuscript.

## Data availability

All data supporting the findings of this study are available within the paper and its Supplementary Information. Additional data are available from the corresponding authors upon reasonable request.

## Figure legend

**Supplementary Figure 1.**
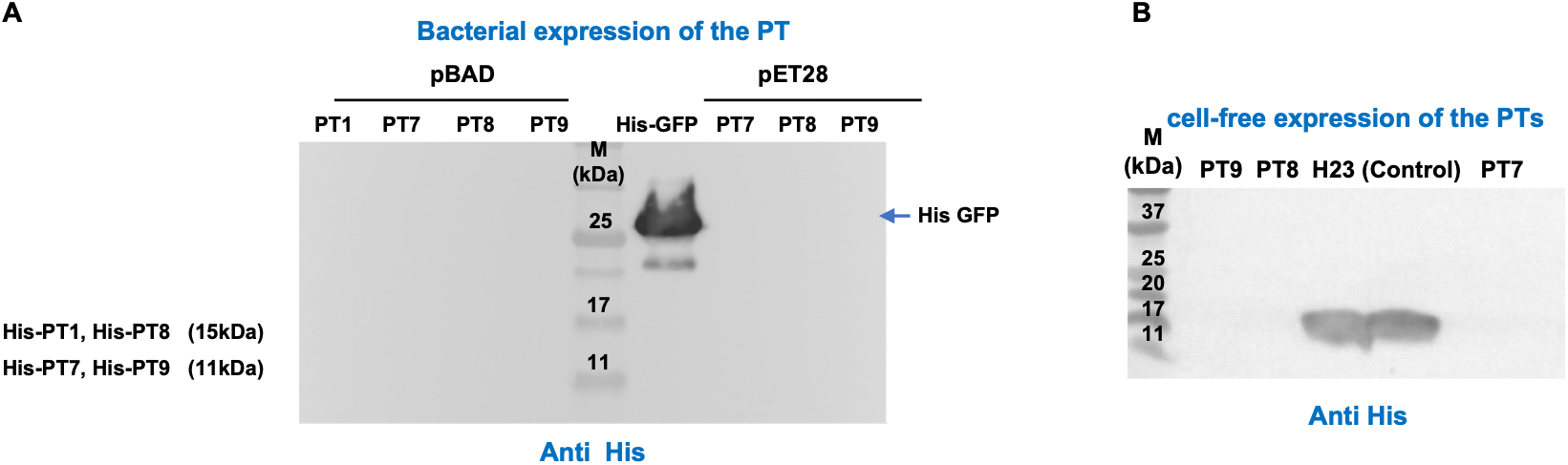
Expression of the PTs using standard recombinant expression approaches. **(A)** Recombinant expression of His-tagged PT7, PT8, and PT9 from pBAD and pET28 vectors. Western blot analysis of small-scale expression cultures showed no detectable PT protein. His-tagged GFP served as a positive control. **(B)** Cell-free expression of His-tagged PT7, PT8, and PT9 using the pIVEX expression system. Western blot analysis revealed no detectable expression of the PT proteins. H23 was included as a positive expression control.

**Supplementary Figure 2.**
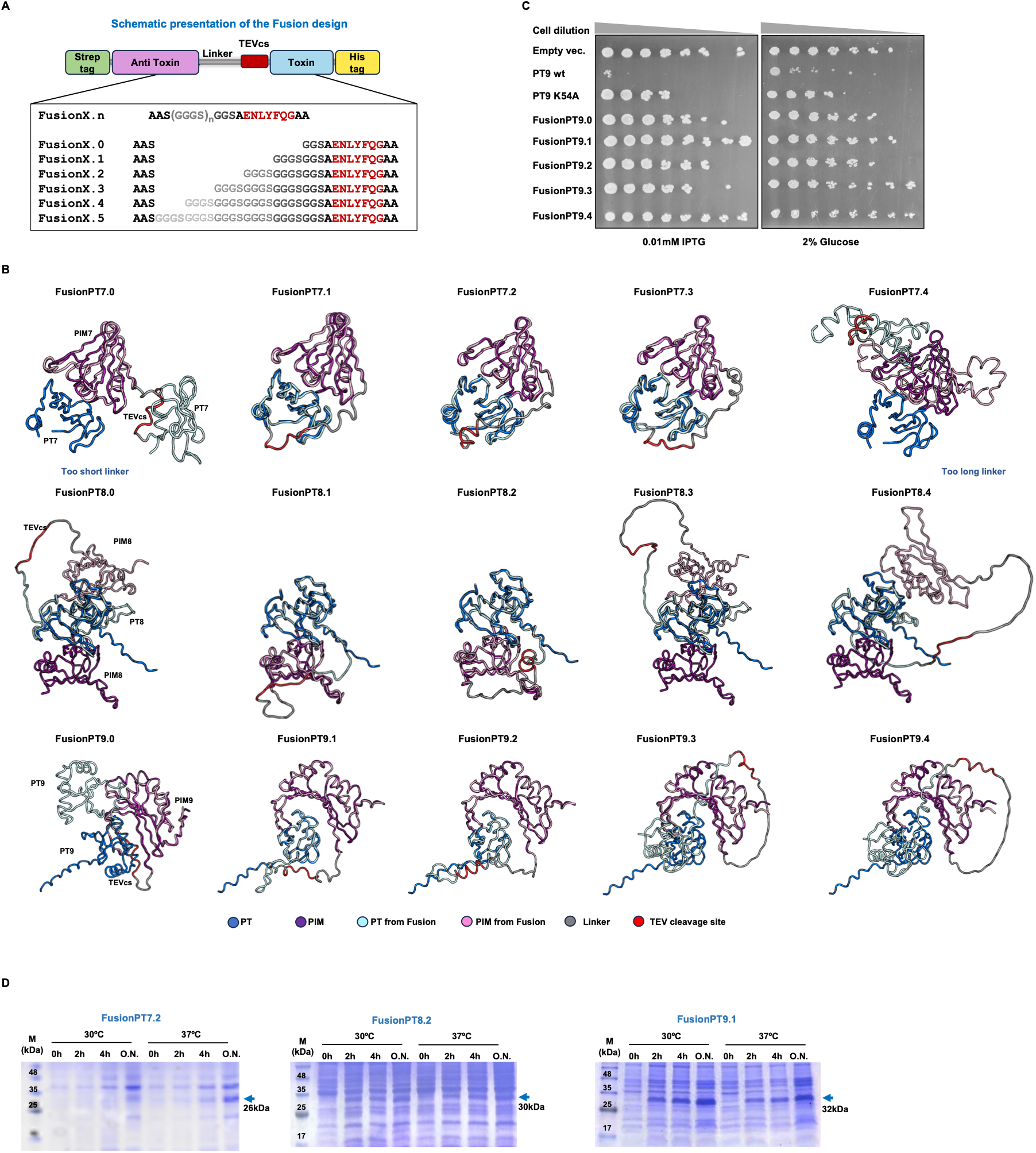
Structure-guided design of FusionPT constructs. **(A)** Schematic representation of FusionPT constructs containing one to five GGGS linker repeats (FusionPT.0–FusionPT.5). (**B**). Structure-guided design of FusionPT7, FusionPT8, and FusionPT9 constructs. Predicted structures of the native non-covalent PT-PIM complexes were superimposed with the corresponding FusionPT constructs containing GGGS linkers of varying lengths to evaluate preservation of the native PT-PIM orientation. **(C)** Effect of linker length on rescue of PT9-mediated toxicity. Drop assays of *E. coli* BL21(DE3) cells expressing FusionPT9.0–FusionPT9.4 under inducing conditions. The catalytically inactive PT9 K54A mutant served as a non-toxic control. **(D)** Optimization of fusion protein expression conditions. SDS-PAGE analysis of small-scale expression of FusionPT7.2, FusionPT8.2, and FusionPT9.1 under different induction temperatures and expression times. Overnight expression produced the highest protein yield.

**Supplementary Figure 3.**
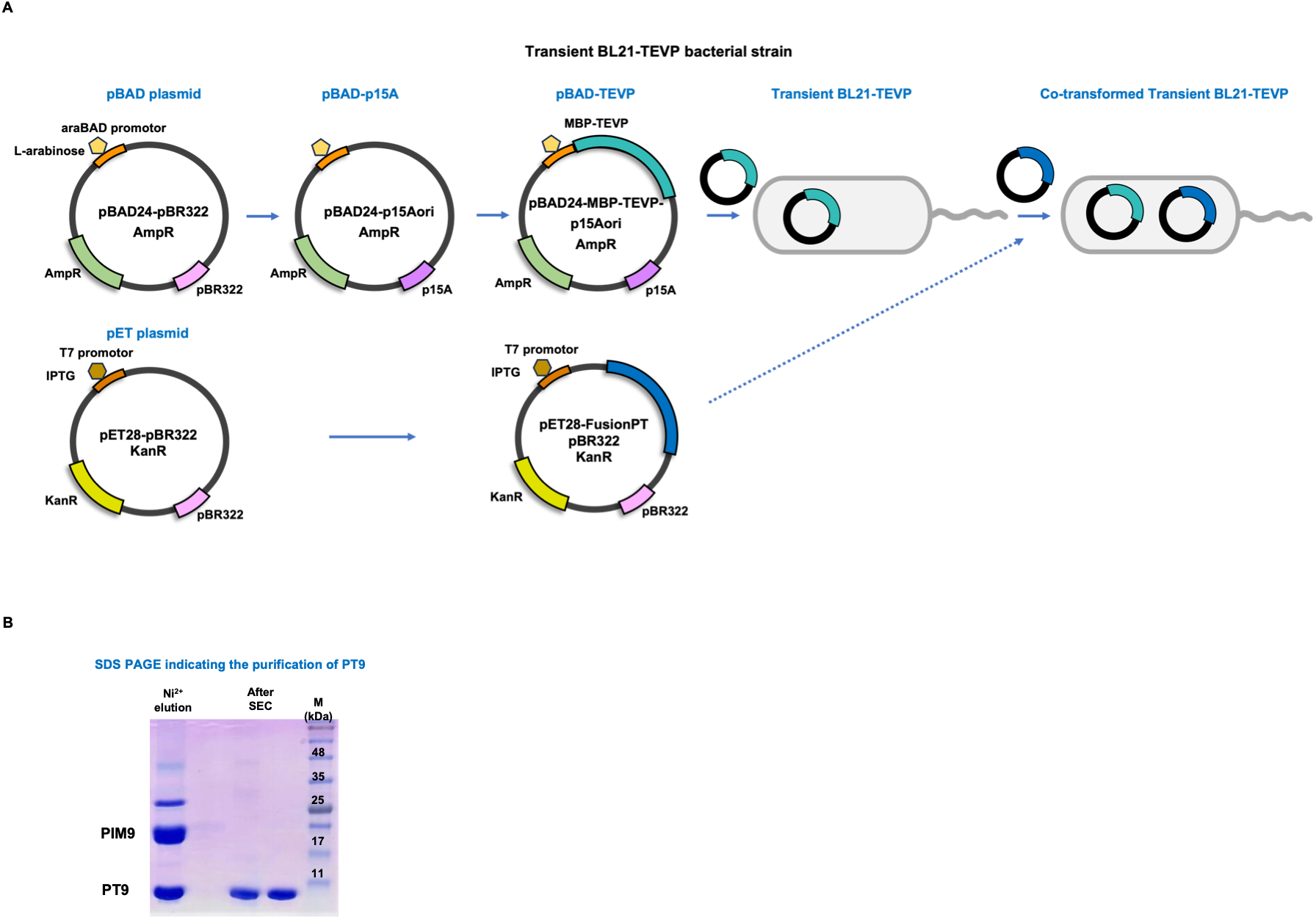
Construction of the transient tBL21-TEVP strain. **(A)** Schematic illustration of the pBAD-MBP-TEVP expression plasmid and generation of the transient BL21(DE3) host used for intracellular TEV-mediated cleavage. (**B**) SDS-PAGE analysis of the purified PT9 following SEC indicating high purity.

**Supplementary Figure 4.**
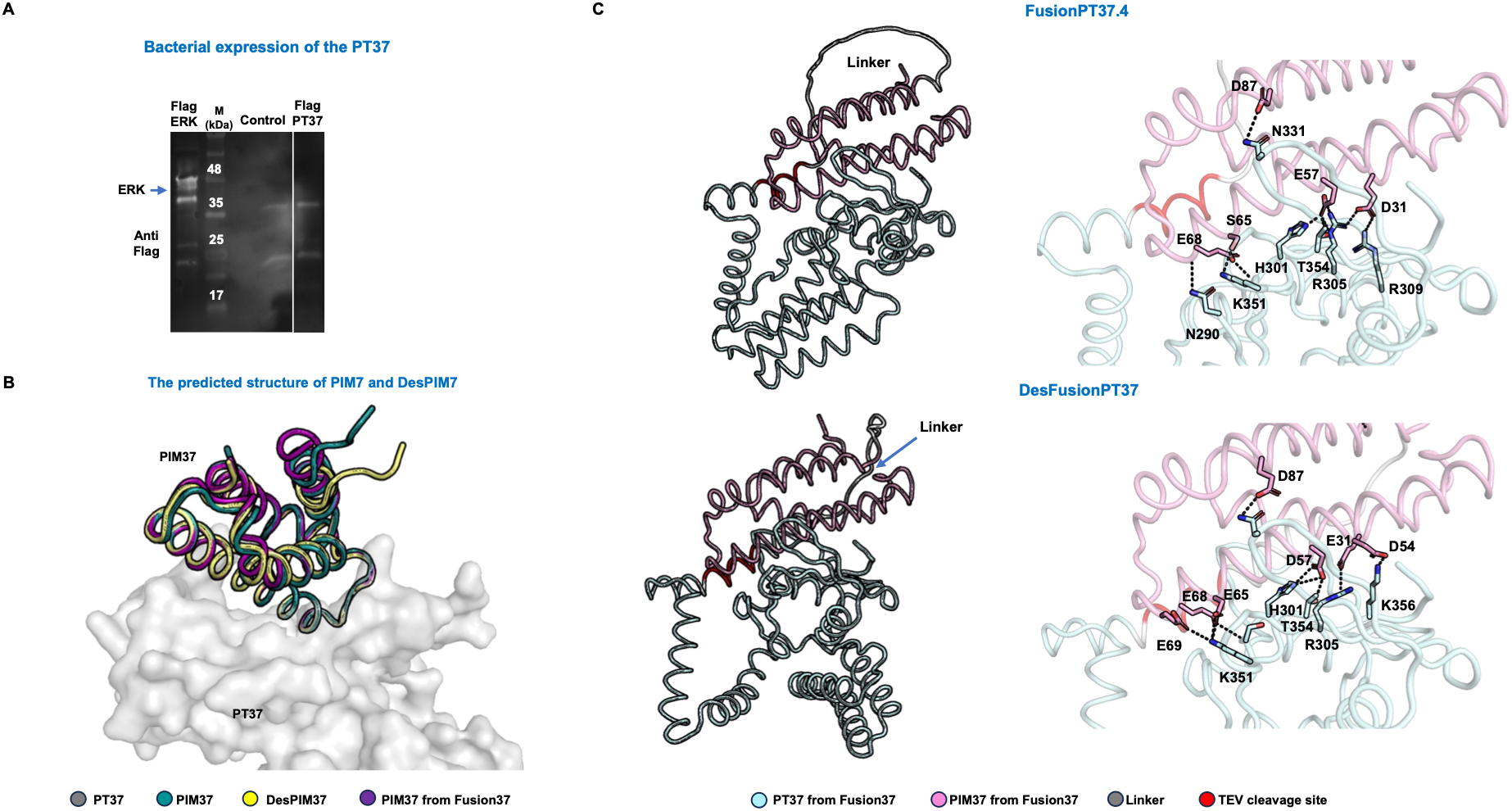
Computational redesign of immunity proteins expands the fusion-based production platform. (A) Western blot analysis of small-scale expression of Flag-tagged PT37 showing no detectable recombinant expression. Cell lysate expressing Flag-tagged ERK was used as a positive control. **(B)** Superposition of the predicted structure of PIM37 and the designed PIM37 on the predicted structure of FusionPT37 indicates a similar conformation of the PIM. **(C)** Structural comparison of the predicted FusionPT37 and DesFusionPT37 constructs. Superposition of the full fusion proteins (left) shows a similar overall conformation of the PT37-PIM37 interface, with local changes in the linker conformation. Comparison of the PT37–PIM37 interfaces (right) indicates that the redesigned construct preserves the overall toxin-immunity interactions.

